# High reproducibility demonstrated in a direct-to-consumer microbiome testing workflow under real-world conditions using fecal reference material

**DOI:** 10.64898/2026.11.04.764294

**Authors:** Julian M. Trachsel, Hannah Sturgeon, David M. Goad, Timothy J. Straub, Ruben A. T. Mars, Cheryl Sew Hoy, Kimberley V. Sukhum

**Affiliations:** Seeding Inc, doing business as Tiny Health, Austin, Texas, USA; Division of Gastroenterology and Hepatology, Mayo Clinic, Rochester, Minnesota, USA

## Abstract

Recent studies have documented substantial variation in microbiome profiles recovered from identical biological samples across direct-to-consumer (DTC) gut microbiome testing providers and, in some cases, within individual providers. These reports underscore the need for transparent, published assessments of analytical performance from the companies offering these services.

To assess analytical reproducibility in our own testing workflow, we processed 10 replicates of a homogenized National Institute of Standards and Technology (NIST) human fecal reference material (RM 8048) through our full sample processing pipeline, from sample collection to data generation. Replicates were shipped from five U.S. sites over a staggered 35-day window. They were processed amongst over 6,000 customer samples, batched for DNA extraction and sequencing based on their arrival in the lab in nine separate sequencing runs, and analyzed using our standard pipeline.

The 10 reference material replicates yielded highly concordant community profiles regardless of shipping time or temperature. Technical variability was minimal (median Bray-Curtis dissimilarity = 0.04) compared with the variation observed in samples from the same Tiny Health customer collected the same day (median Bray-Curtis: 0.08), within one week (0.25), one to two weeks apart (0.28), and between unrelated individuals (0.88). When evaluated using NIST’s Coefficient of Disagreement (COD) metric, our workflow yielded a maximum pairwise COD of 0.045, well below NIST’s published upper bound of 0.073 for this reference material. All three spike-in control strains were successfully recovered from every sample. These results demonstrate the reproducibility of our DTC microbiome testing workflow under real-world conditions, with robust consistency across sample collection, shipping, laboratory processing, and bioinformatics analysis. They also highlight the value of RM 8048 and similar reference materials for evaluating and transparently reporting analytical performance in DTC microbiome testing.

## Introduction

Over the past 25 years, the microbiome has emerged as an important component of health and disease (Huang et al., 2022). As microbiome testing continues to expand across research, clinical, and consumer settings, establishing confidence in the analytical reproducibility of these measurements is increasingly important. For direct-to-consumer (DTC) testing in particular, consumers and researchers need evidence that reported microbiome profiles reflect the underlying sample rather than variability introduced during collection, shipping, laboratory processing, and bioinformatics. This consistency is a prerequisite for accurate results and for valid comparisons across samples, whether between individuals, among research samples, or across longitudinal samples from the same individual.

Recent studies have highlighted methodological variability as a challenge for the reproducibility and comparability of microbiome measurements. Forry et al. (2025) found that microbiome variation as a result of methodological choices could be similar in magnitude to biological differences. In a direct evaluation of DTC gut microbiome testing taking place before Tiny Health had the visibility to be included, Servetas et al. (2026) reported substantial variation within and between DTC providers, with differences between providers sometimes comparable to biological variation between donors. These findings underscore the need to evaluate analytical reproducibility and transparently report performance.

Here, we evaluated the reproducibility of Tiny Health’s end-to-end testing workflow using 10 replicates of a homogenized NIST fecal reference material (RM 8048), shipped from five geographically distant U.S. sites under realistic conditions. We assessed consistency across replicates and concordance with NIST’s independent characterization of the reference material (Jackson et al., 2026). This work demonstrates how fecal reference materials can be used to evaluate a DTC testing workflow, from sample collection through bioinformatics, and provides a model for the transparent reporting of analytical performance called for by recent studies (Servetas et al., 2026).

## Methods

In the same laboratory, 10 Copan 4N6FLOQSwabs® Genetics sample tubes, which are used in the Tiny Health DTC test, were sampled from the same aliquot of the homogenized NIST reference standard (RM 8048, omnivore) and were sent through Tiny Health’s workflow. Two sample tubes per location were first shipped to 5 geographically distant U.S. sites and then from there to a single commercial laboratory over a staggered 35-day window, to simulate the possible delays in collection and shipment of a sample. Total travel time was computed as the interval from collection to receipt at the laboratory. Temperature data during shipment were obtained from the Open-Meteo historical archive and used to estimate a sample’s cumulative exposure to temperatures > 20 °C. At laboratory arrival, the samples were processed for DNA extraction and metagenomic sequencing (∼3 Gbp/sample passing QC) alongside other customer samples, in 9 unique sequencing runs. Resulting data were processed through Tiny Health’s proprietary bioinformatics pipeline; Tiny Health’s Metagenomic Classifier v2 - “THMCv2” (Trachsel et al., 2026).

Reproducibility across the 10 replicates was assessed using Bray-Curtis dissimilarity calculated from species-level relative abundances. These values were put in context against Bray-Curtis dissimilarities calculated from species level taxonomic profiles from pairs of samples from the Tiny Health database in the following categories: pairs from the same-person on the same day, pairs from the same person between 1 and 7 days, pairs from the same-person between 8 and 14 days, and pairs formed from unrelated people. Spearman correlation was used to test the associations between travel time and temperature and dissimilarity. The coefficient of disagreement (COD) was calculated according to NIST SP 260-252 for taxa above 0.0008 relative abundance. Genomes representing the species of the three spiked control strains were present in the taxonomic classification database, and the three spike-ins were identified in the species-level relative abundance profiles.

Genus-level composition profiles were compared between our analysis and NIST’s recently published characterization (Jackson et al., 2026). NIST reports SILVA v138 genus labels while Tiny Health’s current pipeline reports GTDB (Genome Taxonomy Database) r202 (Parks et al., 2020); genus-level abundance across the ten aliquots were translated onto SILVA/NCBI names using a mapping file derived from individual genome assembly accessions and those genomes identities across both taxonomies. Both sides were then canonicalized by stripping SILVA bracket groups and GTDB polyphyly suffixes (Blautia_A → Blautia) and summed within each canonical name. The genera matched this way account for 95.5% of NIST’s published RM 8048 community.

## Results

Recovered community profiles for the 10 samples were nearly identical, with median Bray-Curtis dissimilarities of 0.04 (Figure 1A). This technical variability was substantially lower than the biological variability observed among pairs of Tiny Health same-person customer samples collected on the same day, same-person samples collected either within a one-week or a one- to two-week period, and samples from unrelated individuals (median Bray-Curtis: 0.08, 0.25, 0.28, and 0.88, respectively; Figure 1A). These values are consistent with the range of biological variability reported in another study (Vandeputte et al., 2021), indicating that technical variability was substantially lower than the biological variability observed among customer samples processed in the same batches.

**Figure 1:**
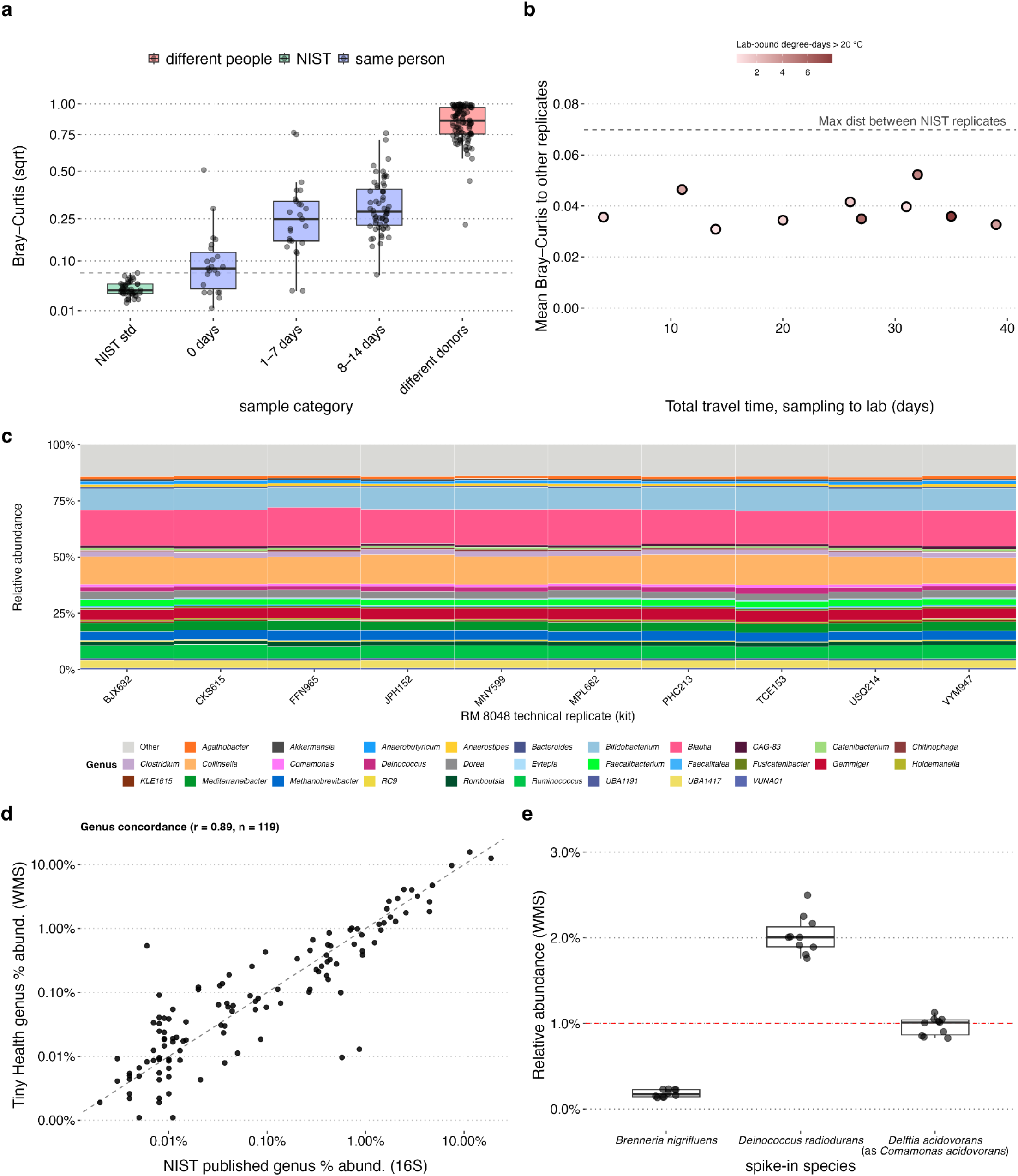
High reproducibility of a microbiome testing workflow using fecal reference material. **(a)** Bray-Curtis dissimilarity among the ten NIST RM 8048 replicates (NIST std), among same-person customer samples grouped by days between collection (0, 1-7, 8-14 days), and among random samples from different individuals (“different donors”). Dashed line indicates maximum NIST replicate dissimilarity. Boxes show median and interquartile range. **(b)** Mean Bray-Curtis dissimilarity of each RM 8048 replicate to the other nine, plotted against total travel time from sample collection to laboratory receipt. Point color indicates cumulative lab-bound thermal exposure above 20°C during shipping. Dashed line indicates maximum NIST replicate dissimilarity. **(c)** Genus-level relative abundance composition of all ten RM 8048 technical replicates, labeled by sample ID. **(d)** Mean genus-level relative abundance concordance between Tiny Health’s shotgun metagenomic sequencing and NIST’s published 16S rRNA-based characterization of RM 8048, on log-transformed axes. Each point represents one genus; dashed line indicates 1:1 concordance. r = 0.89 (Pearson, n = 119 shared genera). **(e)** Relative abundance of the three spiked control strains across all ten replicates, plotted against their 1% target relative abundance (red dashed line). Boxes show median and interquartile range.

This reproducibility held regardless of how long an aliquot traveled. Bray-Curtis dissimilarity between replicates remained low and stable across the full range of travel times examined (4-39 days), and travel time showed no relationship with dissimilarity (Figure 1B, Spearman rho = 0.03, p = 0.95). Estimated cumulative temperature exposure was also not associated with dissimilarity (Figure 1B; Spearman rho = 0.285, p = 0.427). To contextualize our results, we compared our results with NIST’s recently published 16S rRNA-based characterization (Jackson et al., 2026). The coefficient of disagreement (COD, range 0-1, higher values indicate more dissimilarity), a measurement used by NIST to quantify variability across replicates, was calculated for our dataset and compared to the value reported by NIST. The maximum pairwise COD at the genus-level was 0.045, compared to the 0.073 maximal value reported by NIST in their recent characterization of the standard material, further demonstrating the reproducibility of our sample processing pipeline. Genus-level composition profiles across our replicates were similar to those reported by NIST, and were largely composed of *Collinsella, Blautia*, and *Bifidobacterium* (Pearson r = 0.89 on log-transformed relative abundances across 119 genera which were comparable between the 2 different pipelines; Figure 1C, D).

NIST spiked RM 8048 with three organisms as internal controls, each targeted at 1% of cells. All three were detected in all ten of our replicates. *Brenneria nigrifluens* was recovered at approximately one-fifth of target abundance (0.2%, coefficient of variation (CV) 23%), *Deinococcus radiodurans* at approximately twice its target abundance (2.0%, CV 11%), and *Delftia acidovorans* at approximately target abundance (1.0%, CV 11%) (Figure 1E) (Our pipeline reported *Delftia acidovorans* under its previous name, *Comamonas acidovorans*). NIST’s own 16S based characterization of the material shows the same ordering: *Brenneria* (classified as Dickeya by their chosen method) at 0.40%, *Deinococcus* at 1.64%, and *Delftia* at 0.72% (SP 260-252). Because two methods with different biases agree on the order, this suggests the departures from the 1% target reflect a characteristic of the material, whose spike-ins NIST notes are still being characterized, rather than only biases arising from workflows.

Together, these results demonstrate high technical reproducibility across replicates, robustness to variation in shipping conditions, and strong concordance with the independent characterization and internal controls provided by NIST RM 8048.

## Discussion

Technical variability across the ten RM 8048 replicates was substantially lower than the biological variability observed among customer samples, indicating that the testing workflow produces highly consistent results even when samples experience variation in travel time and temperature. The agreement with NIST’s independent characterization was also high despite differences in sequencing approach and taxonomic assignment methodology. These findings further demonstrate how a widely available reference material can provide a common basis for evaluating reliability across microbiome testing providers.

Some disagreement between the two characterizations remained, consistent with differences in sequencing method and reference taxonomic database. Shotgun metagenomics and 16S rRNA sequencing differ in methodological biases and taxonomic resolution, and GTDB and SILVA, the taxonomies used for the respective analyses, can assign organisms differently. Results can therefore differ between providers using different sample processing protocols, sequencing methods, and taxonomies, even when they analyze the same underlying community, and such methodological differences may contribute to the cross-provider disagreement observed in past work (Servetas et al., 2026).

Reconciling differences arising from sample processing, sequencing methods and taxonomies will require broader standardization across the field. In the meantime, we encourage other DTC microbiome testing providers to evaluate their own processes using RM 8048 or a comparable reference material and publish the results. A common landmark gives customers and researchers a measure of reliability independent of any one company’s method or taxonomy, and in doing so, strengthens the field as a whole.

## Author Contributions

JMT, RATM, and KVS conceived the experiment. JMT led the experiment, with all authors contributing to its logistics. JMT performed the analysis and made the figure. JMT and HS wrote the manuscript. All authors reviewed and edited the manuscript.

